# Brain-region-specific changes and dysregulation of activity regulated genes in *Gria3* mutant mice, a genetic animal model of schizophrenia

**DOI:** 10.1101/2024.11.15.623468

**Authors:** Wei-Chao Huang, Ryan Kast, Kira Perzel Mandell, Borislav Dejanovic, Kevin Bonanno, Sameer Aryal, Zohreh Farsi, Jonathan Wilde, Dongqing Wang, Xian Gao, Hasmik Keshishian, Steven A. Carr, Guoping Feng, Morgan Sheng

**Affiliations:** The Stanley Center for Psychiatric Research, The Broad Institute of MIT and Harvard, Cambridge, MA, USA; McGovern Institute for Brain Research and Yang Tan Collective, Massachusetts Institute of Technology, Cambridge, MA, USA; Department of Brain and Cognitive Sciences, Massachusetts Institute of Technology, Cambridge, MA, USA; The Proteomics Platform, The Broad Institute of MIT and Harvard, Cambridge, MA, USA; Capsida Biotherapeutics, Thousand Oaks, CA; Vigil Neuroscience, Cambridge, MA, USA; Emugen Therapeutics, Cambridge, MA, USA

## Abstract

Protein-truncating variants in *GRIA3* (encoding the GluA3/GluR3 subunit of α-amino-3-hydroxy-5-methyl-4-isoxazolepropionic acid (AMPA)-type glutamate receptors) are associated with substantially increased risk of schizophrenia (SCZ). Here we characterized *Gria3* mutant mice carrying a protein-truncating mutation that mimics a SCZ-associated variant. Transcriptomic analysis revealed that activity-regulated genes are downregulated in cortical regions, while immune and glia-related pathways exhibit brain-region-specific changes. The transcriptomic changes in *Gria3* mutant mice are remarkably different from those in *Grin2a* mutant mice, particularly in the prefrontal cortex, even though both encode glutamate receptors and are associated with SCZ risk. Proteomic analysis further demonstrated that loss-of-function of *Gria3* profoundly alters the protein composition of synapses. These findings in a genetic mouse model provide potential insights into the pathophysiological mechanisms underlying SCZ.

## INTRODUCTION

Schizophrenia (SCZ) is a severe mental disorder and a leading cause of disability among young adults^1^. Despite its high prevalence and significant heritability—estimated to be up to 80%^2^— the molecular and neurobiological mechanisms by which genetic factors contribute to SCZ pathophysiology remain unclear. Recent large-scale genomic studies, including genome-wide association studies (GWAS) and exome sequencing, have identified over 280 common variant loci associated with SCZ^3,4^, as well as a sizable number of genes carrying rare loss-of-function (LoF, such as protein truncating) mutations that greatly increase the risk of SCZ even in the heterozygous state^5^. These human genetics discoveries are paving the way for the development of animal models of SCZ that are crucial for investigating potential molecular and circuit mechanisms of the disease and for advancing mechanism-based therapies–a pressing need given the limited efficacy of current antipsychotic treatments.

The glutamate hypothesis is a long standing hypothesis about SCZ^6^ posits that reduced glutamatergic activity is central to the manifestation of SCZ symptoms, a proposition supported originally by pharmacological studies in which N-methyl-D-aspartate receptor (NMDAR) antagonists, such as phencyclidine and ketamine, induce SCZ-like symptoms^7,8^. Human genetics studies have provided further support for this hypothesis. In particular, a recent large-scale meta-analysis of exomes from individuals with SCZ (SCHEMA study) identified ten high confidence genes in which rare protein-truncating variants (PTVs) greatly increase the risk of SCZ^5^. Among these are *GRIN2A* and *the X-linked gene GRIA3*, which respectively encode the GluN2A and GluA3 subunits of N-methyl-D-aspartate (NMDA) and 5-methyl-4-isoxazolepropionic acid (AMPA) receptors, the key mediators of glutamatergic neurotransmission in the central nervous system^5^. Taken together these findings point to a key role of impaired glutamatergic neurotransmission in the etiology of SCZ.

GluA3 functions as one of four pore-forming subunits (GluA1–4) of AMPARs, which are postsynaptic glutamate receptors that mediate rapid synaptic transmission and which drive NMDAR activation at excitatory synapses through depolarization^9^. Widely expressed in the brain, GluA3 is crucial for several physiological processes, including long-term potentiation in hippocampal and cortico-amygdala synapses^10,11^, and synaptic transmission, and activity-dependent plasticity in the auditory system^12^. Thus, *GRIA3* LoF mutations can directly lead to altered synapses, dysfunction of which has consistently been implicated in SCZ based on human genetics^13^ and postmortem gene and protein expression studies^14-16^. Besides SCZ, mutations in *GRIA3* have been linked to neurodevelopmental phenotypes^17^, epilepsy^18,19^, intellectual disability^20,21^, and aggressiveness^22^, further tying GluA3 to cognitive and emotional functions.

Previously, a multi-omics study of *Grin2a* mutant mice revealed brain-region-specific changes in synaptic and other molecular pathways, uncovering potential mechanisms by which *Grin2a* LoF might contribute to SCZ^23^. How LoF of AMPA receptor subunit *GRIA3* affects the brain, however, is poorly understood. To examine the role of *Gria3 in vivo*, we generated a mouse line harboring a *Gria3* PTV identified in the SCHEMA human genetics study. We conducted genome-wide mRNA profiling across multiple brain regions in *Gria3* mutant mice at two ages alongside proteomic analysis of purified synapses. These analyses revealed large scale and region-specific changes in neurons, such as the downregulation of activity-regulated genes (ARGs) in cortex and striatum, and unexpected alterations in glial cells, including the dysregulation of immune and oligodendrocytes-related pathways. Overall, our findings in this *Gria3* genetic model show transcriptomic differences as well as overlap with the *Grin2a* mutant mice and provide further insights into the potential mechanisms underlying SCZ.

## RESULTS

### Dysregulation of activity-regulated and synaptic genes in *Gria3* mutant mice

To gain an unbiased insight into alterations in the brain of *Gria3* mutant mice at both juvenile and young adult stages, we first conducted bulk messenger RNA sequencing (RNA-seq) on *Gria3* hemizygous males (-/y) and their wildtype littermates (+/y) at 1 and 3 months of age (1 and 3mo). The analysis spanned multiple brain regions, including the prefrontal cortex (PFC), hippocampus (HP), somatosensory cortex (SSC), striatum (STR), thalamus (TH), and substantia nigra (SN) (Fig. S1A).

Bulk RNA-seq analysis revealed numerous differentially expressed genes (DEGs, defined as those changing with false discovery rate [FDR]-adjusted P < 0.05) in *Gria3* mutants across all six brain regions and both ages: total number of DEGs across the studied brain regions was 153 (1mo) and 209 (3mo) (total number of unique DEGs: 148 (1mo) and 201 (3mo), Fig. S1A, Table S1). The number of DEGs was greater at 3mo than at 1mo in all regions except the TH, and there was a predominance of downregulated DEGs, particularly in PFC, SSC, STR and TH (Fig. S1A).

To examine the biological processes altered in *Gria3* mutant mice, we conducted gene set enrichment analysis (GSEA) on bulk RNA-seq data for all brain regions and ages. GSEA analysis revealed significant enrichment (FDR < 0.05) of specific molecular pathways and processes (annotated by gene ontology (GO) terms) among the upregulated and downregulated genes (hereafter referred to as upregulated or downregulated pathways/GO terms, respectively).

Notably, the gene-set of ARGs, identified by Tyssowski et al.^24^, was downregulated in most brain regions at both 1 and 3mo, with the most significant changes in cortical regions PFC and SSC (Fig. 1A, Table S2). A subset of ARGs, rapid primary response genes (rapid PRGs; also known as immediate early genes (IEGs)), also showed downregulation in PFC and SSC, more profoundly at 3mo than at 1mo, and to a greater extent in SSC than PFC (Fig. 1A, B). For example, the expression of *Fos, JunB, Arc, Nr4a1* was decreased in the PFC and SSC of *Gria3*^-/y^ males (Fig. 1B). We validated the downregulation of several activity-related DEGs (including *Bdnf*) in SSC using quantitative real-time PCR (qPCR), confirming the robustness of the RNA-seq results and changes in expression of ARGs (Fig. 1C). Besides PFC and SSC, ARGs/IEGs were also downregulated in STR at 3mo, but interestingly, upregulated in HP at the same age (Fig. 1A, B). The expression of ARGs and IEGs can serve as readouts of neuronal activity; therefore, our transcriptomic findings indicate that *Gria3* LoF leads to differential region-specific effects on neuronal activity, with hypoactivity in cortical regions and hyperactivity in HP.

**Figure 1.**
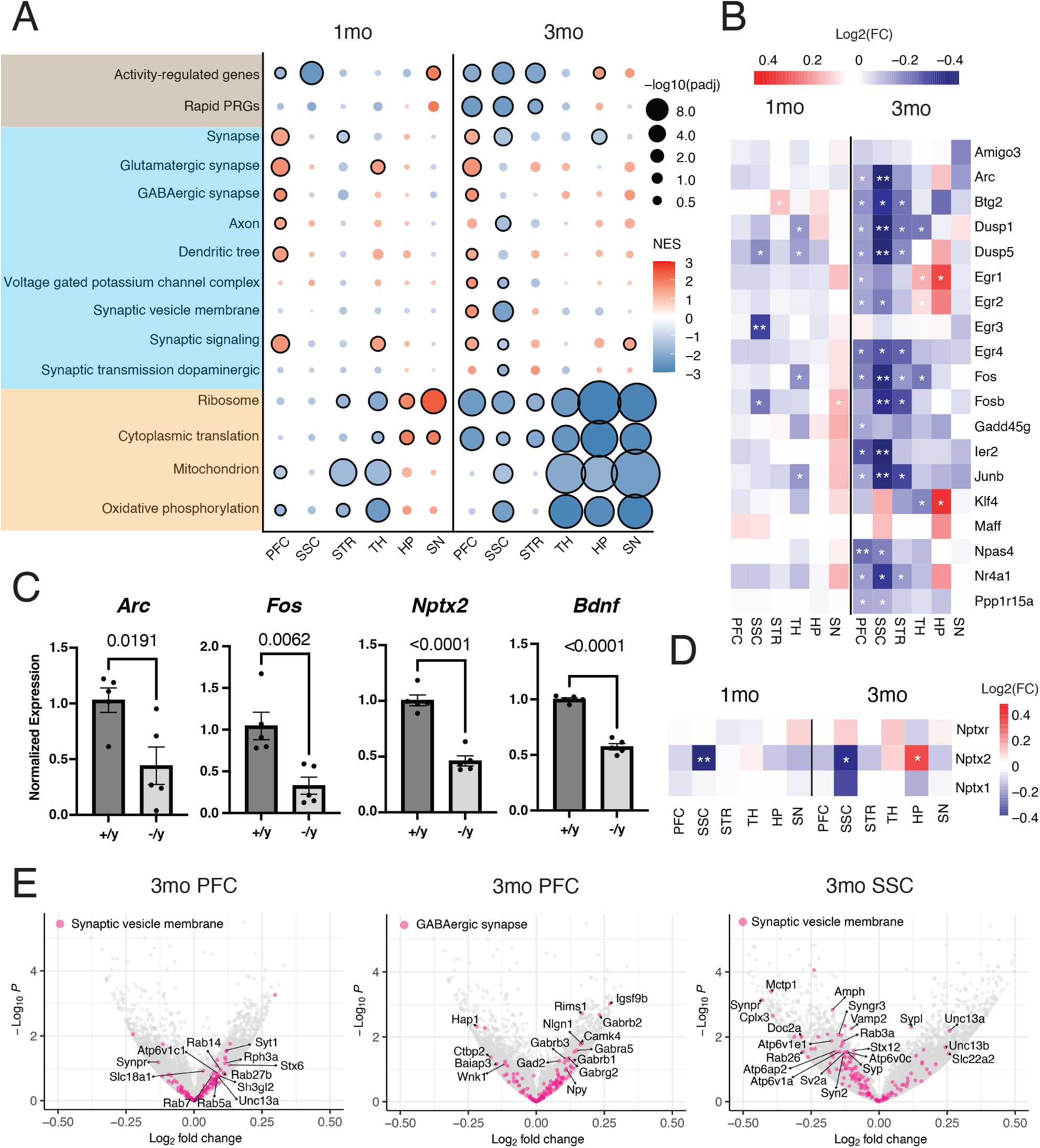
Transcriptomic changes of activity-regulated genes in *Gria3*^-/y^ brain. (A) Gene set enrichment analysis (GSEA) of bulk RNA-seq transcriptomic changes in *Gria3*^-/y^ mice in the indicated brain regions and developmental stages. Statistical significance is represented by the size of the circles and normalized enrichment score (NES) is represented by color scale. Circles with black outlines indicate FDR <0.05. (B) Heatmap showing the log2 fold change (log2FC) values of rapid primary response genes (PRGs) in the bulk RNA-seq data from the indicated brain regions of *Gria3*^-/y^ mice. *: P value < 0.05; **: adjusted P value < 0.05 (C) Comparison of relative mRNA level measured by qRT-PCR (normalized to *Gapdh* mRNA) for indicated genes in the SSC at two ages (3mo: *Arc* and *Fos*; 1mo: *Nptx2* and *Bdnf*) Data are shown as mean +/-standard error (n = 5). Values on each plot are P values computed using two-tailed Student’s t tests. (D) Heatmap showing the log2FC values of *Nptx* genes in the bulk RNA-seq data from the indicated brain regions in *Gria3*^-/y^ mice. *: P value < 0.05; **: adjusted P value < 0.05. (E) Volcano plots of transcriptome changes in *Gria3*^-/y^ mice highlighting genes in the indicated gene sets.

A particularly interesting ARG is Neuronal pentraxin 2 (*Nptx2*; also known as *Np2* or *Narp*)^25^ which plays a role as an extracellular aggregating factor for AMPA receptors^26,27^ and the protein level of which is robustly reduced in the CSF and/or synapse fractions of patients with SCZ^28,29^ and neurodegeneration^30-32^. In *Gria3* mutant mice, the expression of *Nptx2* was reduced in cortical regions at 1 and 3mo, especially in SSC, and elevated in HP at 3mo, correlating with changes in the ARG gene set in these brain regions (Fig. 1D).

Loss of synapses and/or synaptic dysfunction is implicated in SCZ pathophysiology by human genetics and by postmortem brain studies^4,5,16,33^. Consistent with an impact on synapses, GSEA of *Gria3*^-/y^ brain showed significant enrichment of synapse-related GO terms in up- and down-regulated genes (Fig. 1A). In the PFC, synaptic GO terms were upregulated at both 1 and 3mo, whereas in the SSC, these terms were downregulated at 3mo (Fig. 1A). For instance, the GO terms “Synaptic vesicle membrane” and “GABAergic synapse” were upregulated in the 3mo PFC, along with the expression change of associated genes such as *Syt1* (Synaptic vesicle membrane) and *Gabrb2* (GABAergic synapse) (Fig. 1E). In contrast, the gene set of the “Synaptic vesicle membrane” GO term was downregulated in the 3mo SSC, showing an opposite trend compared to the PFC (Fig. 1A, E). GSEA using refined SynGO terms^34^ revealed transcriptomic changes in both presynaptic and postsynaptic processes in *Gria3* mutants (Fig. S1B). Notably, “translation at presynapse” and “translation at postsynapse” GO terms were strongly downregulated across all brain regions at 3mo (Fig. S1B), which mirrors the downregulation of ribosome/translation-related GOs at this age (Fig. 1A). In the PFC, GO terms related to pre- and postsynaptic organization were upregulated, while in the SSC, presynaptic organization was downregulated with minimal change in postsynaptic terms (Fig. S1B).

### Brain-region-specific changes in multiple molecular pathways in 3mo *Gria3* mutant mice

Excessive activation of innate immunity and microglia are thought to contribute to pathogenesis of SCZ and other CNS disorders^32,35-38^. It was therefore interesting that GSEA on bulk RNA-seq data revealed significant alterations in immune-related GO terms at 3mo (Fig. 2A). The GO terms such as “innate immune response”, “reactive astrocytes”, and “inflammatory response”, were downregulated in the PFC, STR, and TH, while being upregulated in the SSC and HP (Fig. 2A, B, Table S2).

**Figure 2.**
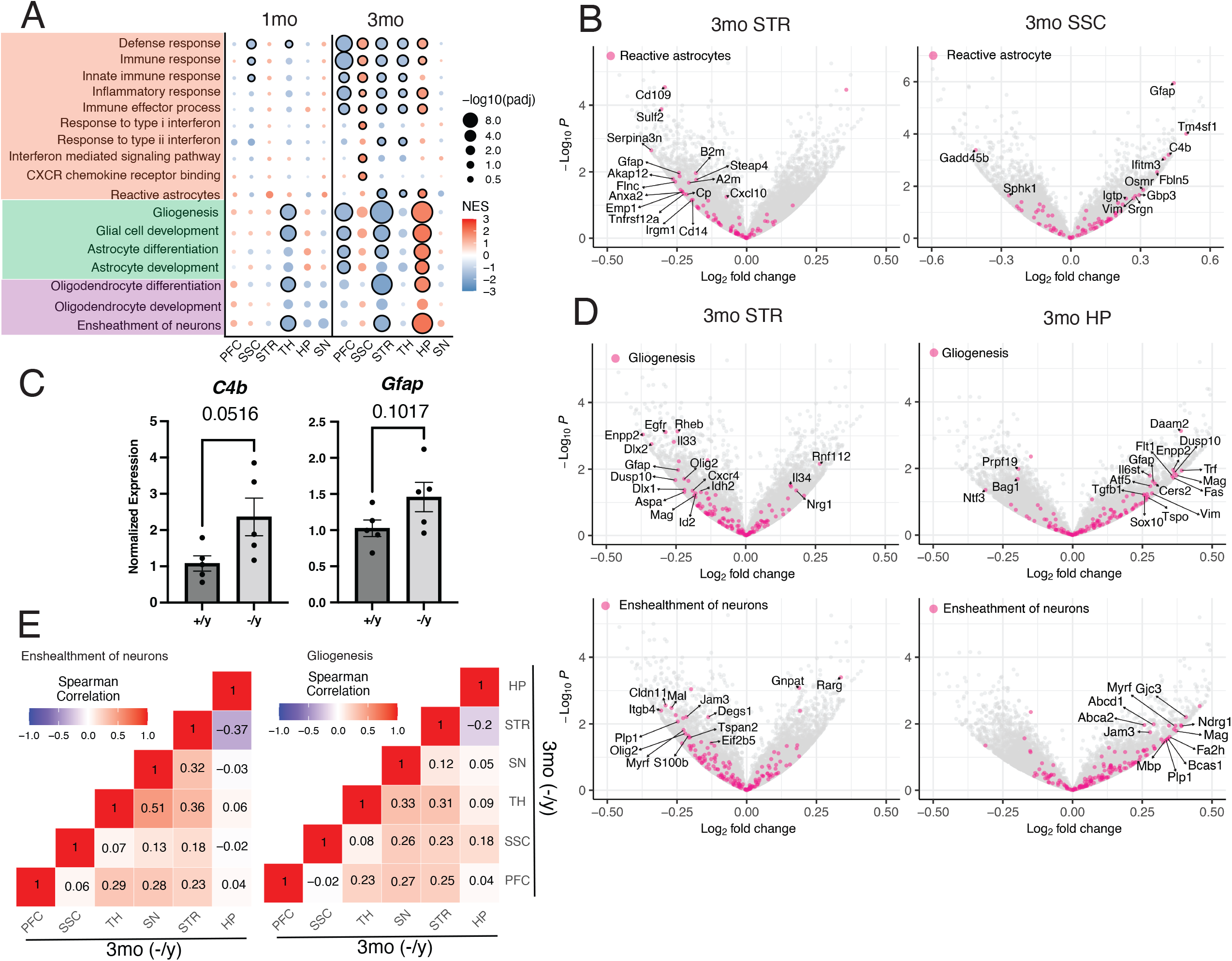
Brain-region-specific changes in immune- and glia-related pathways in 3mo *Gria3* mutant mice. (A) GSEA results of bulk RNA-seq transcriptomic changes in *Gria3*^-/y^ mice at 1 and 3mo in the indicated brain regions and ages. (B & D) Volcano plots of transcriptome changes in *Gria3*^-/y^ mice highlighting genes in the indicated gene sets. (C) qRT-PCR results (normalized to *Gapdh* mRNA) for indicated genes in the 3mo SSC. Data are shown as mean +/-standard error (n = 5). Values on each plot are P values computed using two-tailed Student’s t test. (E) Correlation of RNA-seq results across brain regions. Spearman’s correlation was performed on the log2FC for each pair of brain regions with indicated gene sets. Number in each square represents Spearman’s correlation coefficient.

In the 3mo SSC, where there is downregulation of ARGs and synapse GO terms including *Nptx2* (Fig 1A, D), we noted upregulation of genes associated with reactive astrocytes (Fig. 2A, B). Of particular note, the expression of *C4b* (the single mouse homolog of the human *C4A* and *C4B* complement factor genes, and expressed mainly by astrocytes), was upregulated in the SSC of 3mo *Gria3* mutant mice (Fig. 2B). *C4* has been identified as a copy number genetic variant associated with SCZ^36^, and its overexpression in mice causes excessive synapse loss and SCZ-related phenotypes^35,37,39^. NPTX2 has also been reported to regulate complement activity in the brain^32^. Another gene that is associated with activated astrocytes, *Gfap*, was also robustly elevated in SSC at 3mo (Fig. 2B) We validated the increased expression of *C4b*, along with *Gfap*, an astrocytic activation marker^40^, using qPCR (Fig. 2C).

In addition to immune-related pathways, we observed brain-region-specific alterations in astrocyte- and oligodendrocyte-related pathways (Fig. 2A). For instance, the glial cell markers such as *Gfap* (astrocytes) and *Mag* (oligodendrocytes) were upregulated in HP but downregulated in STR at 3mo (Fig. 2D). We further examined whether similar genes were driving these GSEA changes by examining the correlation across brain regions in the log2FCs (fold changes) of genes constituting these sets. For example, in the GO terms “ensheathment of neurons” and “gliogenesis,” the PFC, TH, and STR showed consistently positive correlations with each other (Spearman’s r = 0.23–0.36), whereas the HP displayed either no correlation or negative correlations with these regions (Spearman’s r = -0.37 to 0.09) (Fig. 2E). This suggests that the similar GSEA changes in PFC, TH and STR might be driven by an overlapping set of genes.

Several pathways exhibited a similar trend, where the direction of changes in the HP and SSC was opposite to that observed in the STR, TH, and PFC at 3mo. For example, GO terms such as “extracellular matrix structural constituent” and “Wnt signaling” were upregulated in the HP and SSC but downregulated in the other three brain regions (Fig. S2A, B). While “RNA splicing” GO term was upregulated in the STR and TH but downregulated in the HP and SSC (Fig. S2A). Overall, these results indicate that *Gria3* LoF induces profound, brain-region-specific transcriptome alterations at 3mo, tending to affect in opposite directions HP and SSC versus PFC, STR and TH. Similar patterns of brain-region-specific transcriptome alterations were observed in *Grin2a* mutant mice, further emphasizing the opposing direction of several GO terms between the SSC and HP versus the STR and TH^23^.

### Sex-specific effects of *Gria3* LoF on multiple brain regions

Especially because previous studies suggested *GRIA3* as a risk gene for SCZ in females^41^, we performed RNA-seq on the same six brain regions of *Gria3* heterozygous (+/-) and homozygous (-/-) female mice at 1mo. The total number of DEGs in *Gria3*^*-/-*^ females (146) was comparable to those in 1mo *Gria3*^-/y^ males (153) (Fig. S1A, S3A, Table S1). The number of DEGs in *Gria3*^-/-^ mice was similar to that in *Gria3*^+/-^ across most brain regions (Fig. S3A). Unlike 1mo *Gria3*^-/y^ males, where the TH had the greatest number of DEGs, the HP exhibited the highest number in female mutants (Fig. S1A, S3A). A transcriptome-wide comparison showed that log2FC values of all individual genes were moderately well-correlated between *Gria3*^+/-^ and *Gria3*^-/-^ female mutants in each brain region studied (Fig. S3B; Spearman r = 0.37–0.62); however, the correlation of transcriptomic change between *Gria3*^-/-^ females and *Gria3*^-/y^ males was weak or non-existent (Fig. S3B). These findings suggest that transcriptome profiles in the same brain regions considerably overlap between *Gria3*^*+/-*^ *and Gria3*^*-/-*^ females, whereas there are notable differences in transcriptomic change between male and female *Gria3* mutants at 1mo.

We performed GSEA on bulk RNA-seq data to assess alterations in biological processes in female mice (Fig. S3C, Table S2). Similar to the findings in males, we observed downregulation of ARGs/IEGs in the SSC and PFC of both *Gria3*^*+/-*^ and *Gria3*^*-/-*^ females (Fig. S3C, S3D). Synaptic GO terms were predominantly downregulated in the PFC but upregulated in the HP in both *Gria3*^*+/-*^ and *Gria3*^*-/-*^ females, consistent with changes in ARG expression (Fig. S3C). The synapse-related GSEA changes in PFC appear to be different than those in the male mutant (Fig. 1A).

We next examined glial cell- and immune-related pathways in 1mo female mice. These pathways were downregulated in the PFC and HP in both mutant genotypes (+/-& -/-) (Fig. S3C), which contrasts with 1mo male mutants that showed minimal changes in these pathways (Fig. 2A). GO terms such as “extracellular matrix structural constituent” and “Wnt protein binding,” which showed the same direction of change as glial/immune-related GO terms in 3mo *Gria3*^-/y^ males (Fig. 2A, S2B), also exhibited the same patterns in females, being downregulated in both the PFC and HP (Fig. S3C, S3E).

Overall, we found that ARGs/IEGs were predominantly downregulated in the SSC of *Gria3* mutant mice of both sexes (Fig. 1A, 1B, S3C, S3D). However, comparisons of the GSEA results between male and female *Gria3* mutant mice revealed several sex-specific effects of *Gria3* LoF. For instance, synapse-related GO terms were upregulated in the PFC of 1mo males but downregulated in females. Additionally, brain-region-specific effects observed in 1mo female mutants (e.g., downregulation of glial/immune-related GO terms in the PFC and HP) were not present in 1mo mutant males, although distinct brain-region-specific alterations in these pathways were observed in 3mo males.

### Differences in transcriptome profiles between *Grin2a* and *Gria3* mutant mice

Our previous research identified brain-region-specific changes and evidence of dysregulation of dopamine signaling in *Grin2a* mutant mice^23^. Given that both *Grin2a* and *Gria3* are glutamate receptor genes implicated in SCZ by the SCHEMA study, we sought to compare their LoF transcriptomic phenotypes. We compared transcriptomes of *Gria3*^-/y^ males with *Grin2a*^-/-^ males, as hemizygous *Gria3* LoF is functionally similar to a homozygous knockout.

Surprisingly, GSEA showed that the gene set of ARGs exhibited opposite directions of change between *Gria3*^-/y^ and *Grin2a*^-/-^ mice across most brain regions at 1 and 3mo (Fig. S4A). For instance, in the PFC and SSC, ARGs were upregulated in *Grin2a*^-/-^ mutants but downregulated in *Gria3*^-/y^ mutants (Fig. S4A). A comparison of log2FC of ARGs in the 3mo PFC and SSC showed a moderate inverse correlation between the two mutants (Fig. S4B). For example, *Nptx2* expression decreased in *Gria3*^-/y^ SSC but increased in *Grin2a*^-/-^ SSC (Fig. S4C).

In the PFC, multiple pathways also showed opposite GSEA changes between *Gria3*^-/y^ and *Grin2a*^-/-^ mutants (Fig. S4A). Synaptic GO terms were upregulated in *Gria3*^-/y^ but downregulated in *Grin2a*^-/-^ mutants. Changes in gene sets related to glutamatergic and GABAergic synapses were negatively correlated between the *Gria3* and *Grin2a* mutants (Fig. S4D, Spearman r ≈ -0.34). Additionally, immune-related GO terms in the 3mo PFC were upregulated in *Grin2a* mutants but downregulated in *Gria3* mutants (Spearman r: innate immune response: -0.289; inflammatory response: -0.225).

### Single nucleus RNA-seq changes in neuronal and non-neuronal cells in *Gria3* mutant mice

To investigate transcriptomic changes at the cell-type specific level, we performed single-nucleus RNA-seq (snRNA-seq) in the PFC of *Gria3*^-/y^ males at 1 and 3mo. Additionally, the TH (1mo) and STR (3mo) of *Gria3*^-/y^ males were studied with snRNA-seq as these brain regions showed large changes in bulk RNA-seq at the respective ages. No significant differences in cell type proportions were observed between *Gria3* mutants and wildtype littermates (Table S3).

GSEA of snRNA-seq data revealed that synapse-related GO terms were upregulated in both excitatory and inhibitory neurons in the PFC at 3mo (Fig. 3A, Table S4). This finding is consistent with the results from bulk RNA-seq data in the 3mo PFC (Fig. 1A). However, there were no significant changes in synapse-related GO terms in PFC neurons at 1mo (Fig. 3A). To determine which specific neuronal subtypes exhibited these alterations at 3mo, we conducted further analysis and found that synapse-related GO terms were significantly and most prominently upregulated in layer 6 corticothalamic (L6 CT) neurons, along with downregulation of ARGs (Fig. 3A, B).

**Figure 3.**
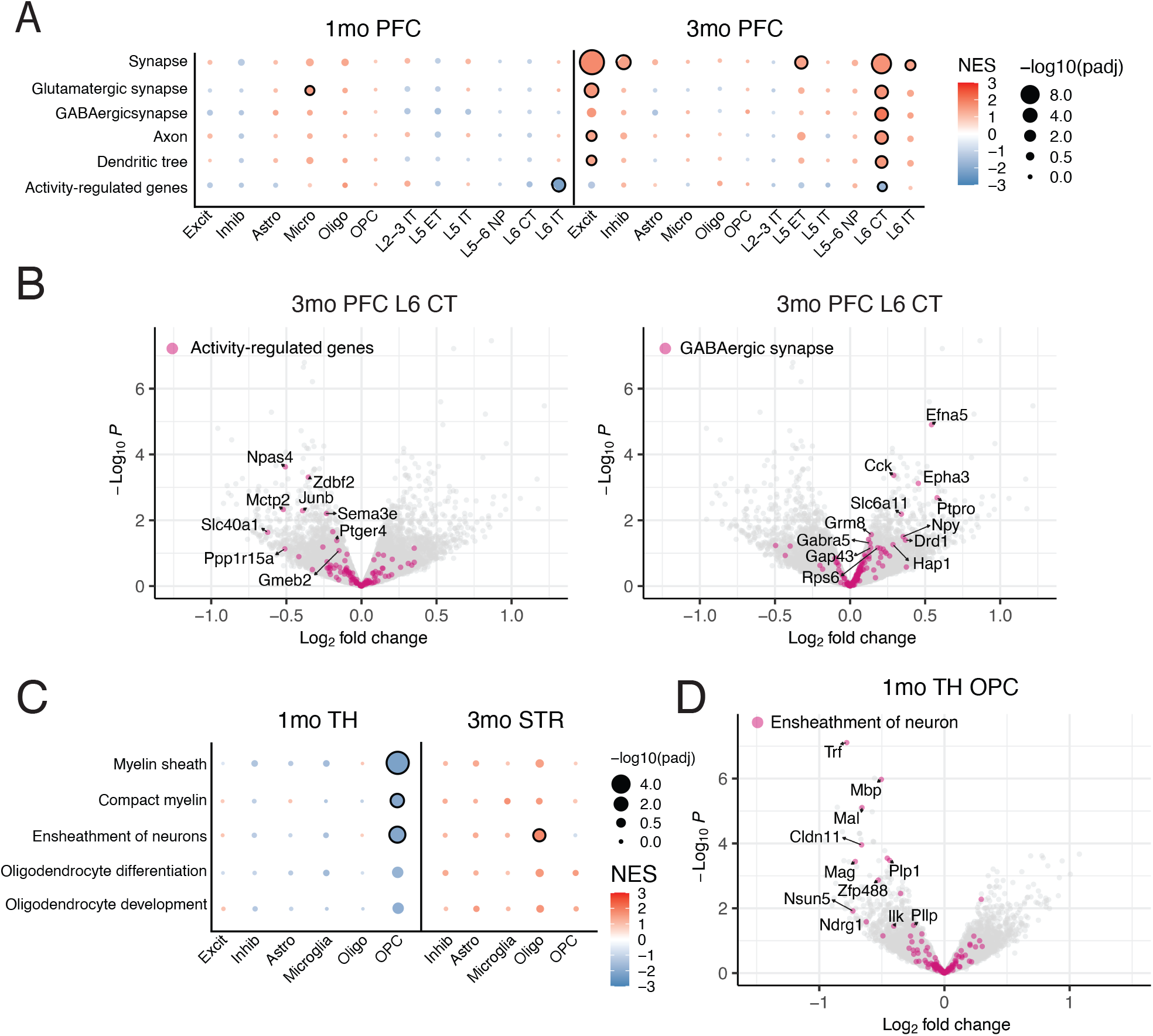
Transcriptomic changes in neuronal and non-neuronal cells in *Gria3* mutant mice. (A) GSEA results of single nucleus RNA-seq (snRNA-seq) transcriptomic changes in Gria3^-/y^ mice at 1 and 3mo in the indicated brain regions and ages. Excit, excitatory neuron; Inhib, inhibitory neuron; Astro, astrocyte; Micro, microglia; Oligo, oligodendrocyte; OPC, oligodendrocyte progenitor cells; L2–L6, layers 2–6; IT, intratelencephalic; ET, extratelencephalic; NP: near-projecting; CT, corticothalamic neurons (B) Volcano plots of transcriptome changes in the 3mo PFC L6 CT of *Gria3*^-/y^ mice highlighting genes in the indicated gene sets (C) GSEA results of snRNA-seq transcriptomic changes in the 1mo TH and 3mo STR of *Gria3*^-/y^ mice. (D) Volcano plots of transcriptome changes of snRNA-seq in *Gria3*^-/y^ mice highlighting genes in the indicated gene sets.

AMPA receptors are expressed in oligodendrocyte precursor cells (OPCs) and mature oligodendrocytes, where they are required for oligodendrocyte functions^42^, such as cell proliferation^43^ and myelination^44^. In GSEA of bulk RNA-seq data from *Gria3*^*-/y*^ mutants, we noticed downregulation of several myelin- and oligodendrocyte-related GO terms in the TH at 1mo and in the STR at 3mo (Fig. 2A). To identify the specific cell types driving these changes, we performed snRNA-seq on the 1mo TH and 3mo STR of *Gria3*^*-/y*^ mice and conducted GSEA analysis. The results showed that myelin- and oligodendrocyte-related GO terms were primarily downregulated in OPCs of 1mo TH of Gria3^-/y^ mutants (Fig. 3C, D). No cell type of STR showed downregulation of myelin or oligodendrocyte-related terms by snRNA-seq analysis (Fig. 3C).

### Changes in synapse proteomics of *Gria3* mutant mice

To investigate the impact of *Gria3* LoF on synapses at the protein level, we performed quantitative mass spectrometry (MS)-based proteomics on synaptic fractions purified from the cerebral cortex of *Gria3*^-/y^ males at 1 and 3mo. GSEA of the synaptic proteomics data revealed significant downregulation of GO terms related to synapses, including synapse, postsynapse, presynapse, excitatory synapse, and inhibitory synapse, in *Gria3*^-/y^ mice at 1mo (Fig. 4A, Table S6). At 3mo, several synapse-related GO terms, such as synapse, presynapse, and postsynapse, also remained downregulated (Fig. 4A). Remarkably, the excitatory synapse GO term shifted from downregulation at 1mo to upregulation at 3mo, with several NMDA receptor subunits (GRIN1, GRIN2A, GRIN2B), the AMPA receptor subunit (GRIA1), and their associated scaffolding proteins (DLG4/PSD-95, SHANK1, HOMER1) showing increased abundance at 3mo (Fig. 4A, B). Accordingly, the term “glutamate receptor activity” was upregulated at 3mo. These changes likely reflect a compensatory response to the loss of AMPA receptor subunit GluA3 from synapses.

**Figure 4.**
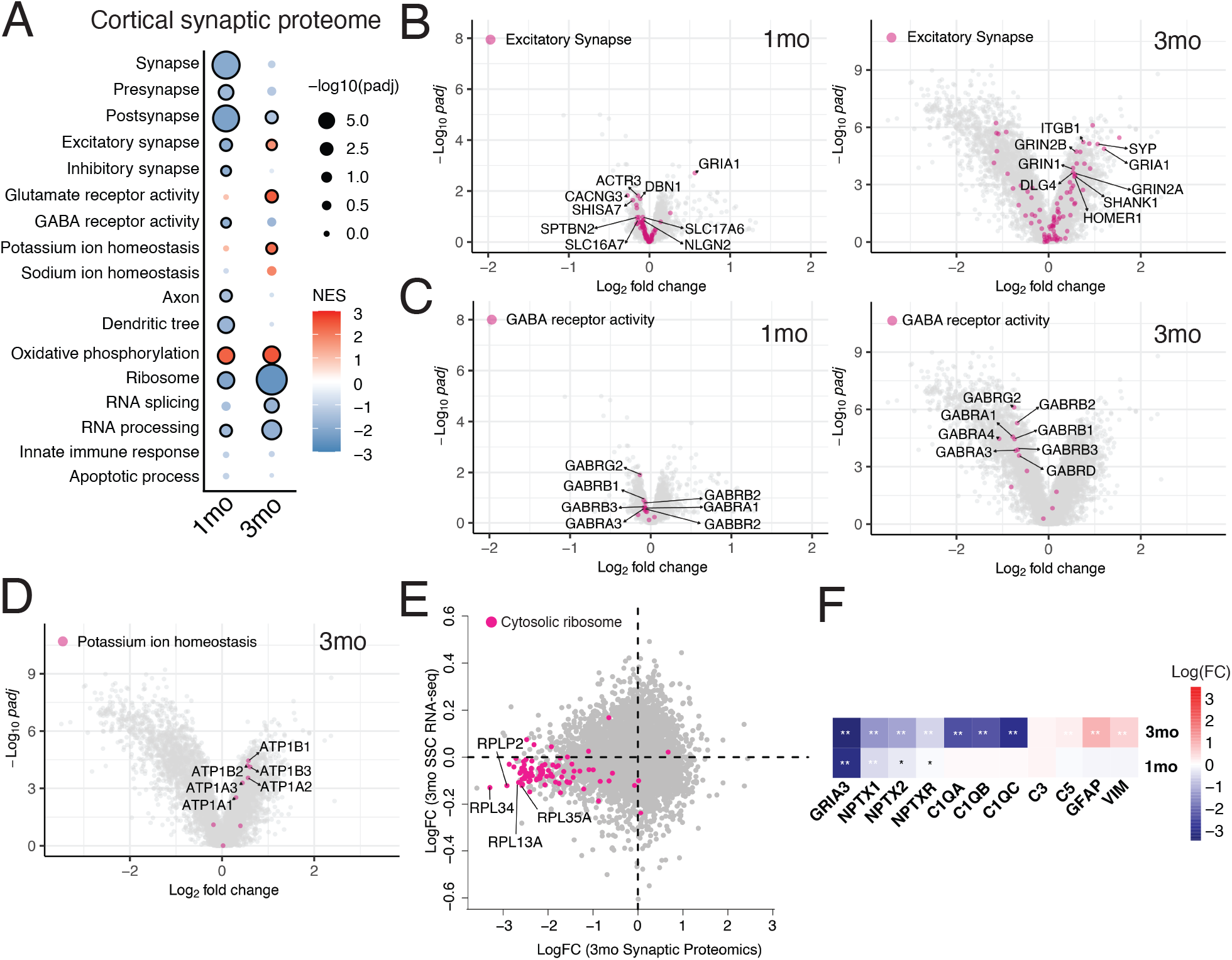
Alteration of synaptic proteome in *Gria3*^-/y^ cortex. (A) GSEA results of the synaptic proteome of 1 & 3mo *Gria3*^-/y^ cortex. (B, C, D) Volcano plots of proteome changes in *Gria3*^-/y^ mice highlighting genes in the indicated gene sets (E) Scatter plot with the transcriptomic changes in the 3mo SSC versus proteomic changes in the 3mo cortex of *Gria3*^-/y^ mutants, highlighting the cytosolic ribosome gene set. (F) Heatmap showing the log2FC values of indicated proteins in the synaptic proteomics data from the indicated ages in *Gria3*^-/y^ mice. *: P value < 0.05; **: adjusted P value < 0.05.

In contrast to the increased levels of glutamate receptors (which mediate excitatory synaptic transmission), GABA receptor subunits (e.g., GABRA1, GABRB1, GABRB2, GABRB3, which mediate inhibitory transmission), were predominantly reduced in the synapse proteome of *Gria3*^-/y^ mutants (Fig. 4B). We hypothesize that this change might again be a compensatory mechanism for the loss of *Gria3*, an excitatory receptor subunit. Additionally, GO terms related to potassium and sodium homeostasis were upregulated at 3mo, with increased levels of Na^+^/K^+^ ATPase pump subunits (e.g., ATP1B1 and ATP1B3) at the synapse (Fig. 4A, D). These results show that GRIA3 deficiency profoundly affects the protein composition of synapses, but in quite different ways at 1 and 3mo.

Changes in ribosomal proteins (decreased), RNA processing proteins (decreased) and in oxidative phosphorylation / mitochondria proteins (increased) were observed in the synaptic proteomes of *Gria3*^-/y^ mice at 3mo and, to a lesser extent, at 1mo (Fig. 4A). More than 90% of downregulated ribosomal proteins in the cortical synaptic proteome of 3mo *Gria3*^-/y^ mice were also downregulated at the mRNA level in the 3mo SSC (Fig. 4E), suggesting that transcriptional change might be contributing, at least in part, to the abundance of ribosomes at synapses.

Finally, we noted that the levels of NPTX proteins (especially NPTX1, NPTX2) were reduced in the purified synapses of *Gria3*^-/y^ mutants (Fig. 4F), aligning with RNA-seq results from the SSC (Fig. 1D). C1Q subunits, known as NPTX2 binding proteins^32^, and also as opsonins for synapse elimination by microglia^32,45^ were also strongly decreased at 3mo, while markers of astrocyte activation, such as GFAP and VIM, were upregulated (Fig. 4F).

## DISCUSSION

In this study, we performed transcriptomic and proteomic analyses on *Gria3* mutant mice, an animal model of SCZ based on human genetics, to capture a comprehensive view of gene expression and protein level changes in the brain caused by *Gria3* LoF. We also compared transcriptomic profiles between *Gria3* and *Grin2a* mutant mice, two rare-variant large-effect risk genes that encode glutamate receptor subunits.

AMPA receptors play a crucial role in rapid synaptic transmission, making them key mediators of neuronal activation. Changes in ARGs/IEGs expression, a readout of neuronal activity, may occur when AMPA receptor function is impaired. Previous studies have shown that AMPA receptors regulate the expression of some IEGs, such as *Arc*^46,47^. Our bulk RNA-seq analysis revealed that *Gria3* LoF reduces the expression of many IEGs in different brain regions, consistent with the function of *Gria3* as an important component of excitatory synaptic transmission.

We observed that *Gria3* LoF has brain-region-specific effects on the brain transcriptome, similar to *Grin2a* mutants, but these two mutations affect different regions in contrasting ways. Surprisingly, the LoF effects of these two SCZ-linked genes caused largely opposite changes in the expression of activity-regulated and synapse-related genes, which may support the contrasting phenotypes observed in previous studies of *Gria3* and *Grin2a* mutant mice. *Gria3* knockout mice exhibited hypoactivity in the open field test^48^, whereas *Grin2a* mutant mice demonstrated hyperactivity^49^. Additionally, *Gria3* knockout mice showed reduced EEG power during NREM sleep^48^, while *Grin2a* mutants showed increased NREM EEG power^23,49^. Further studies are needed to examine the phenotypes of *Gria3* mutant mice and compare them with those of *Grin2a* mutant mice. Nonetheless, our findings here—admittedly on animal models of a complex human disease--presage that the neurobiological mechanisms that underlie SCZ are likely to be extremely heterogeneous.

Our synaptic proteomic analysis revealed significant alterations in synaptic proteins in *Gria3* mutant mice, including changes in subunits of glutamate receptors such as GRIA1 (GluA1) and GRIN2B (GluN2B). These findings are consistent with previous studies indicating that *Gria3* knockout can influence the expression of glutamate receptor subunits. For example, double knockout of GluA1 and GluA3 reduces GRIA2 (GluA2) levels but increases GRIN2A and GRIN2B levels in the brain^50^. Similarly, loss of GluA3 expression decreases synaptic GluA2 and increases synaptic GluA4 subunits in the mid cochlea^51^. Our synaptic proteomics results further support the notion that loss of GluA3 affects the level of AMPA and NMDA receptor subunits and provide a comprehensive view of GluA3 loss-induced alterations in synaptic proteins. A new insight from our work is that the synaptic changes resulting from *Gria3* LoF changes drastically over time.

In conclusion, our findings reveal transcriptomic and proteomic changes caused by SCZ-related *Gria3* PTVs across different developmental stages and brain regions, offering new insights into the molecular mechanisms underlying the clinical phenotypes associated with *Gria3* PTV mutations.

## ACKNOWLEDGEMENTS

We thank Xiao-Man Liu and Chuhan Geng for valuable input for this manuscript; Sean K. Simmons and Joshua Z. Levin for their advice on transcriptomic analysis.

## AUTHOR CONTRIBUTIONS

W.-C. H. and M.S. conceived and designed the experiments. The manuscript was written by W.-C. H. and M.S., with inputs from all authors. W.-C. H. performed sample and library preparations for bulk and snRNA-seq and conducted data analysis and visualization. R.K., J.W., D.W., and X.G. generated and maintained the mouse line under the guidance of G.F. K.P.M. processed the transcriptome data, including quality control, alignment, cell type clustering, annotation, and differential expression analysis. B.D. purified synapse fractions for proteomics. K.B. performed liquid chromatography (LC)-MS/MS for synapse proteomics under the guidance of H.K. and S.A.C. S.A. and Z.F. assisted with data analysis.

## DECLARATION OF INTERESTS

M.S. is cofounder and scientific advisory board (SAB) member of Neumora Therapeutics and serves on the SAB of Biogen, Proximity Therapeutics and Illimis Therapeutics. S.A.C. is a member of the SAB of Kymera, PTM BioLabs, Seer, and PrognomIQ.

## METHODS

### Preparation of injection mixtures

The double stranded DNA donor (dsDNA) for targeted knock-in of the *Gria3*^R210X^ mutation was generated by PCR, cloned into pBluescriptII, and mutated using the Q5 site-directed mutagenesis kit (New England Biolabs, E0554S). Modified sgRNAs targeting *Gria3* were synthesized by Synthego (see Table for sgRNA and donor sequences). All injection mixtures were prepared in a final volume of 50µL according to the following protocol. Using RNase-free water, reagents, and consumables, sgRNA (final concentration 0.61µM) and ultrapure Tris-HCl, pH 7.39 (final concentration 10mM, ThermoFisher) were mixed and EnGen Cas9 NLS, *S. pyogenes* (New England Biolabs, M0646T) was added to a final concentration of 30ng/µL. The mixtures were incubated at 37°C for 15 minutes before adding dsDNA donor (10ng/µL). Injection mixtures were stored on ice and briefly heated to 37°C prior to injection.

### Natural mating for zygotic injections

Three days prior to microinjection, female mice (4-5 weeks old, C57BL/6NTac) were superovulated by IP injection of pregnant mare serum gonadotropins (PMS, Prospec Bio, Cat # HOR-272; 5 IU/mouse) and human chorionic gonadotropin (HCG, Prospec Bio, Cat # HOR-250; 5 IU/mouse, 47 hours after PMS injection) and then paired with males. Donor females were sacrificed by cervical dislocation at day 0.5pcd and zygotes were collected into 0.1% hyaluronidase, EmbryoMax® FHM HEPES buffered medium (FHM, Sigma-Aldrich, MR-025-D). Zygotes were washed in drops of FHM and cumulus cells were removed. Zygotes were cultured EmbryoMax® KSOM-AA (Sigma-Aldrich, MR-107-D) for one hour and then used for microinjection.

### Zygotic microinjections

All microinjections were performed using a Narishige Micromanipulator, Nikon Eclipse TE2000-S microscope, and Eppendorf 5242 microinjector. Individual zygotes were injected with 1-2pL of injection mixture using an “automatic” injection mode set according to needle size and adjusted for clear increase in pronuclear volume. Following injections, cells were cultured in EmbryoMax® KSOM-AA (Sigma-Aldrich, MR-107-D) for 24 hours and then surgically implanted into pseudopregnant CD-1 females (Charles River Laboratories, Strain Code 022) and allowed to develop normally until natural birth.

### Breeding and husbandry

The *Gria3*^R210X^ founder line was backcrossed to C57Bl/6J for a minimum of 6 generations prior to being used for experiments. Experimental mice were the progeny of crosses involving *Gria3* heterozygous females (*Gria3*^+/-^) with wildtype (*Gria3*^+/y^) or *Gria3* hemizygous males (*Gria3*^-/y^). All mice had ad libitum access to food and water, and were housed under a standard 12:12 hour light/dark cycle (lights on at 07:00, lights off at 19:00). Mice were housed in cages of 2-5 mice with their littermates.

### Brian perfusion and dissection

Detailed procedure of brain perfusion and dissection was described previously^23^. In brief, Mice were anesthetized with 3% isoflurane and transcardially perfused using ice-cold Hank’s Balanced Salt Solution (HBSS, Life Technologies) to remove blood from the brain. The brains were then frozen in liquid nitrogen vapor and stored at -80°C. Brain dissections were performed in a cryostat (Leica), with regions including medial prefrontal cortex, dorsal hippocampus, thalamus, somatosensory cortex, dorsal striatum, and substantia nigra meticulously isolated using a precooled biopsy punch and microscalpel. Brain regions were identified using the Allen Brain Atlas, and dissected tissues were stored at -80°C in 1.5ml Eppendorf tubes.

### Sample and library preparation for transcriptomic analysis

We followed the protocols published previously to prepare samples and libraries^23^. RNA was extracted from micro-dissected brain tissue using the RNeasy Mini Kit (Qiagen), with RNA concentration and integrity assessed by NanoDrop and Agilent 2100 Bioanalyzer, ensuring RNA integrity numbers (RIN) above 7 for all samples. The purified RNA was stored at -80°C until library preparation for bulk RNA-seq. Libraries were prepared using the TruSeq Stranded mRNA Kit (Illumina) from 200 ng of total RNA per sample, and the resulting cDNA libraries were quantified using High Sensitivity DNA chips on the Agilent 2100 Bioanalyzer. A 10 nM normalized library pool was sequenced on a NovaSeq S2 (Illumina) with 50 base-pair read lengths.

For snRNA-seq library preparation, nuclei were extracted from dissected brain tissue using a gentle, detergent-based dissociation method, as outlined in a published protocol on protocols.io^52^. The extracted nuclei were enriched via fluorescence-activated cell sorting (FACS) using a Sony SH800 sorter at the Flow Cytometry Core facility of the Broad Institute. RNA from the nuclei was captured with the Chromium v3.1 kit (10x Genomics), and library preparation followed the manufacturer’s instructions. A 10 nM normalized library was pooled, and sequencing was performed on a NovaSeq S2 (Illumina) with paired-end, dual indexing format, with reads of 28 and 75 bases for reads 1 and 2, respectively.

### Purification of synaptic fraction for mass spectrometry (MS) and quantitative MS analysis

The methods for this study were adapted from our previous papers^23,28,53,54^. Briefly, frozen cortices were collected following the method mentioned above in a cryostat using microscalpel. Cortex tissue was thawed and homogenized in ice-cold homogenization buffer (5 mM HEPES pH 7.4, 1 mM MgCl_2_, 0.5 mM CaCl_2_, with phosphatase and protease inhibitors). The homogenate was centrifuged at 1,400 g for 10 minutes at 4°C, and the supernatant was further centrifuged at 13,800 g for 10 minutes. The resulting pellet was resuspended in 0.32 M sucrose, 6 mM Tris-HCl (pH 7.5), layered onto a discontinuous sucrose gradient (0.85 M, 1 M, 1.2 M), and ultracentrifuged at 82,500 g for 2 hours. The synaptosome fraction at the 1 M/1.2 M sucrose interface was collected, mixed with 1% Triton X-100, incubated on ice for 15 minutes, and ultracentrifuged at 32,800 g for 20 minutes to obtain the synapse (postsynaptic density) fraction. This fraction was resuspended in 1% SDS, protein concentration was measured using a BCA assay, and the sample was stored at -80°C.

For MS/MS analysis, eighteen synaptic fractions from wild-type and *Gria3*^-/y^ mouse cortices were analyzed using tandem mass tag (TMT) isobaric labeling for quantification (1mo wildtype n = 4; 3mo wildtype n = 4; 1mo *Gria3*^-/y^ n = 5; 3mo *Gria3*^-/y^ n = 5).

Proteins in the synapse fraction samples (in 1% SDS) were reduced with 5 mM dithiothreitol and alkylated using 10 mM iodoacetamide at room temperature. The denatured and alkylated proteins were processed using S-Trap technology (Protifi) as per the manufacturer’s instructions, with contaminants removed via centrifugation. Digestion was performed on-column using a 1:25 enzyme-to-substrate ratio of each Lys-C and Trypsin overnight at room temperature.

After digestion, 20 μg of each sample was labeled with TMT18 reagent, with samples from each group randomly assigned to channels within the TMT18 plex. Label incorporation was confirmed to exceed 95%, after which reactions were quenched with 5% hydroxylamine and pooled. The TMT18-labeled peptides were desalted using a 50 mg tC18 SepPak cartridge and fractionated by high pH reversed-phase chromatography on a 4.6mm x 250 mm Zorbax 300 Extend-C18 column (Agilent). One-minute fractions were collected throughout the elution and concatenated into 12 fractions for LC-MS/MS analysis.

One microgram of each proteome fraction was analyzed on a Exploris 480 QE mass spectrometer (Thermo Fisher Scientific) coupled to a Easy-nLC 1200 system (Thermo Fisher Scientific). Samples were separated using 0.1% Formic acid / 3% Acetonitrile as buffer A and 0.1% Formic acid / 90% Acetonitrile as buffer B on a 25cm 75um ID picofrit column packed in-house with Reprosil C18-AQ 1.9 mm beads (Dr Maisch GmbH) with a 110 min gradient consisting of 2-6% B in 1 min, 6-20% B in 62 min, 20-30% B for 22 min, 30-60% B in 9 min, 60-90% B for 1 min followed by a hold at 90% B for 5 min. The MS method consisted of a full MS scan at 60,000 resolution and a normalized AGC target of 100% and maximum inject time of 25 ms from 350-1800 m/z followed by MS2 scans collected at 45,000 resolution with a normalized AGC target of 200% with a maximum injection time of 50 ms and a dynamic exclusion of 15 seconds. Precursor fit filter was used with a threshold set to 50% and window of 1.2 m/z. The isolation window used for MS2 acquisition was 0.7 m/z and 20 most abundant precursor ions were fragmented with a normalized collision energy (NCE) of 32 optimized for TMT18 data collection.

Mass spectra were analyzed using Spectrum Mill MS Proteomics Software (Broad Institute) with a mouse database from Uniprot.org downloaded on 04/07/2021 containing 55734 entries. Search parameters included: ESI Q Exactive HCD v4-35-20 scoring, parent and fragment mass tolerance of 20 ppm, 40% minimum matched peak intensity, trypsin allow P enzyme specificity with up to four missed cleavages and calculate reversed database scores enabled. Fixed modifications were carbamidomethylation at cysteine. TMT labeling was required at lysine, but peptide N termini could be labeled or unlabeled. Allowed variable modifications were protein N-terminal acetylation, oxidized methionine, pyroglutamic acid, and pyro carbamidomethyl cysteine. Protein quantification was achieved by taking the ratio of TMT reporter ions for each sample over the TMT reporter ion for the median of all channels. TMT18 reporter ion intensities were corrected for isotopic impurities in the Spectrum Mill protein/peptide summary module using the afRICA correction method which implements determinant calculations according to Cramer’s Rule and correction factors obtained from the reagent manufacturer’s certificate of analysis (https://www.thermofisher.com/order/catalog/product/90406) for lot numbers VH310017 and WG333575.

After performing median-MAD normalization, a moderated two-sample t-test was applied to the datasets to compare wildtype and *Gria3*^-/y^ sample groups at 1 and 3mo. A comprehensive list of DEPs for 1 and 3mo *Gria3*^-/y^ cortical synapse fraction samples is provided in Table S5.

### Bulk RNA-seq analysis

Raw FASTQ files from each sequencing experiment were aligned to a reference using genome FASTA and transcriptome GTF files extracted from the CellRanger mm10 reference for alignment and quantification. Quantification was performed with Salmon^55^ (version 1.7.0) using parameters -l A --posBias --seqBias --gcBias --validateMapping, with a Salmon reference index that included genomic decoys. Quality metrics were generated using STAR^56^ (version 2.7.10a) and Picard tools (version 2.26.7, Broad Institute). Differential expression analysis was conducted between heterozygous/hemizygous/homozygous mutant and wild-type mice for each brain region and age using the DESeq2 R package^57^ (version 1.34). The Salmon output was loaded using tximport^58^ (version 1.22), and genes with at least 10 counts across all samples in each experiment were included in the analysis. Log2 fold change shrinkage was applied using the “normal” shrinkage estimator in DESeq2.

### Single-nucleus RNA-seq analysis

FASTQs were generated from raw BCL files and aligned to the mm10 mouse reference genome using the Cell Ranger pipeline^59^ (v6.1.2). The --chemistry=SC3Pv3 flag was applied during Cell Ranger count, with the --expect-cells parameter set based on nucleus counts from microscopy. Replicates within each experiment (brain region and age) were downsampled using Cell Ranger aggr. Seurat^60,61^ (v4.0.3) was used for further processing and analysis of Unique Molecular Identifier (UMI) counts, with nuclei expressing fewer than 500 genes removed. The remaining nuclei were log-normalized and scaled by a factor of 10,000, followed by scaling with ScaleData for dimensional reduction. Linear dimension reduction was performed by RunPCA on variable genes, followed by clustering using FindNeighbors (with 20 dimensions) and FindClusters, and visualization via Uniform Manifold Approximation and Projection (UMAP) using 20 dimensions. Doublets were identified and removed using Scrublet^62^ (v0.2.3), and small clusters predominantly labeled as doublets were discarded. Major cell types were annotated based on marker gene expression, and neuronal nuclei were re-clustered and further classified into subtypes using Azimuth^60^ (v0.4.6). Cell type proportions were summarized, and statistical differences were evaluated with speckle’s propeller function^63^ (v0.0.3). Differential expression analysis was performed using a pseudobulking approach, where counts were summed for each gene across all cells of each cell type per replicate. The pseudobulk counts were processed with edgeR^64^ (v3.36.0), and lowly expressed genes were filtered using the filterByExpr function. Surrogate Variable Analysis^65^ (SVA, v3.42.0) was employed to identify significant unknown latent sources of noise, which were included as covariates in the differential expression model. The likelihood ratio test was used to compare differential expression between mutant and wild-type replicates.

### Gene set enrichment analysis (GSEA)

GSEA was performed using the negative log10 of nominal P values, multiplied by the sign of log2FC, derived from bulk RNA-seq, snRNA-seq, and proteomics data. The R package fGSEA^66^ v1.3.0 package was used for analysis with gene sets from the Molecular Signature Database^67^ (M5 v2023.1), SynGO^34^, and gene sets curated from the literature (see Table S7). For proteins with multiple isoforms, the isoform with the highest spectral count was used for GSEA. Mouse gene symbols from transcriptomics and proteomics data were mapped to their human homologs using Ensembl’s BioMart^68^, allowing GSEA to be performed with human gene symbols. All significant Gene Ontology (GO) terms (FDR < 0.05) across datasets are listed in the supplementary tables.

### Quantitative realtime-PCR

The method for RNA extraction was mentioned above. Reverse transcription was carried out with the iScript kit (Bio-Rad) following the manufacturer’s instructions. Quantitative real-time PCR was performed using PowerTrack SYBR Green Master Mix (ThermoFisher) on a BioRad CFX384 Real-Time PCR Detection System. The primer information is provided in Table S7. Statistical comparisons for quantification were made using a two-tailed t-test.

## FIGURE LEGENDS

**Figure S1. Widespread transcriptomic changes in different brain regions in *Gria3*^-/y^ mice**

(A) Number of DEGs in the indicated brain region and age in *Gria3*^-/y^ mutants. The size of the circles represents the number of differentially expressed genes (DEGs), defined as those that differ from wildtype with an adjusted P value <0.05. The color of the circle represents the direction of the expression changes. (B) GSEA of bulk RNA-seq with SynGO terms in *Gria3*^-/y^ mice from the indicated brain regions and ages. Statistical significance is represented by the size of the circles and normalized enrichment score (NES) is represented by color scale. Circles with black outlines indicate FDR <0.05.

**Figure S2. Brain-region-specific changes in multiple molecular pathways in 3mo *Gria3* mutant mice**

(A) GSEA results of bulk RNA-seq transcriptomic changes in *Gria3*^-/y^ mice at 1 and 3mo in the indicated brain regions and ages. (B & D) Volcano plots of transcriptome changes in *Gria3*^-/y^ mice highlighting genes in the indicated gene sets.

**Figure S3. Transcriptomic changes in the brain of *Gria3* mutant females**

(A) Number of DEGs in the indicated brain region and age in *Gria3* mutant females. The size of the circles represents the number of DEGs, defined as those that differ from wildtype with an adjusted P value <0.05. The color of the circle represents the direction of the expression changes (B) Correlation of RNA-seq results across brain regions. Spearman’s correlation was performed on the log2FC for each pair of brain regions. Number represents Spearman’s correlation coefficient. (C) GSEA results of bulk RNA-seq transcriptomic changes in *Gria3*^+/-^ *and Gria3*^-/-^ females in the indicated brain regions and ages. (D) Heatmap showing the log2FC values of rapid PRGs in the bulk RNA-seq data from the indicated brain regions in *Gria3* mutant females. *: P value < 0.05; **: adjusted P value < 0.05. (E) Volcano plots of transcriptome changes in *Gria3* mutant females highlighting genes in the indicated gene sets.

**Figure S4. Comparison on the transcriptome profiles between *Gria3*^-/y^ and *Grin2a*^-/-^ mice**

(A) GSEA of bulk RNA-seq transcriptomic changes in *Gria3*^-/y^ and *Grin2a*^-/-^ mice. Most of GSEA results of *Gria3* mutant mice have been shown in the previous figures. (B & D) Scatter plots with transcriptomic changes in the 3mo SSC or 1mo PFC of *Gria3*^-/y^ mice versus *Grin2a*^-/-^ mice, highlighting the indicated gene sets. Spearman’s r correlation values of the indicated gene sets are labeled on the plots. (C) Heatmap showing the log2FC values of *Nptx* genes in the bulk RNA-seq data from the indicated brain regions of *Gria3*^-/y^ and *Grin2a*^-/-^ mice. *: P value < 0.05; **: adjusted P value < 0.05. The GSEA and gene expression results of *Gria3* mutant mice shown in this figure were presented in previous figures of this manuscript, while those of *Grin2a* mutant mice have been published previously. We re-presented these results for comparison.

