## Supplementary figures and images for "Brain-region-specific changes and dysregulation of activity regulated genes in *Gria3* mutant mice, a genetic animal model of schizophrenia"

### Supplemental Figures

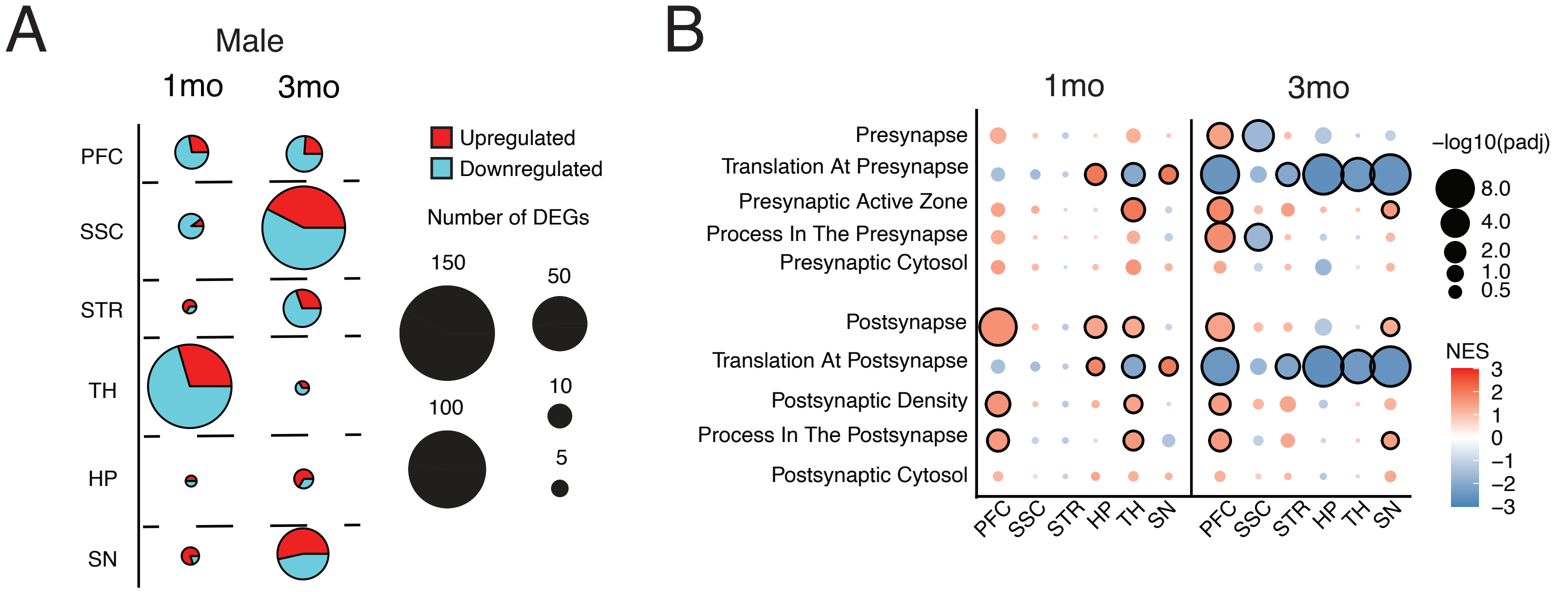

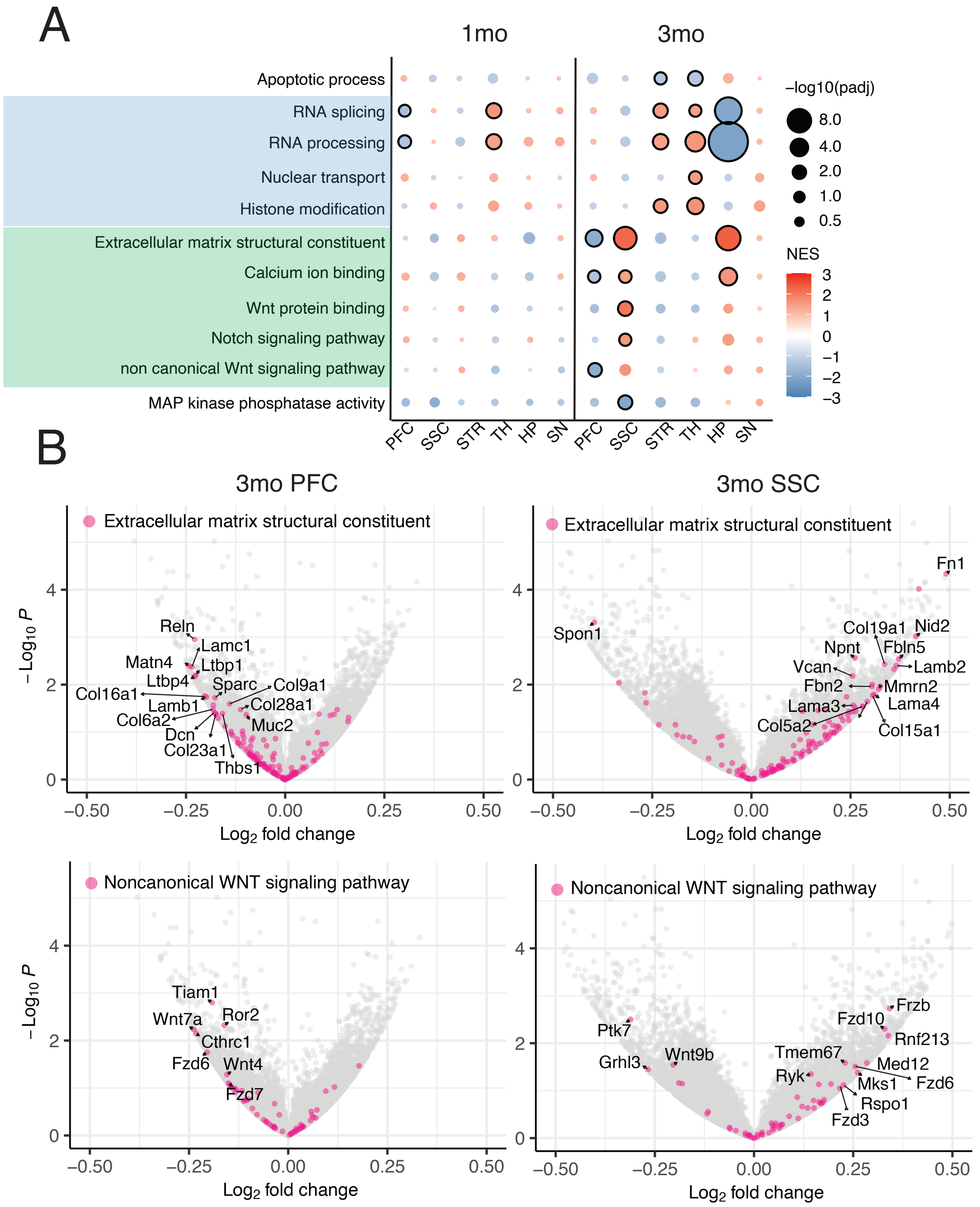

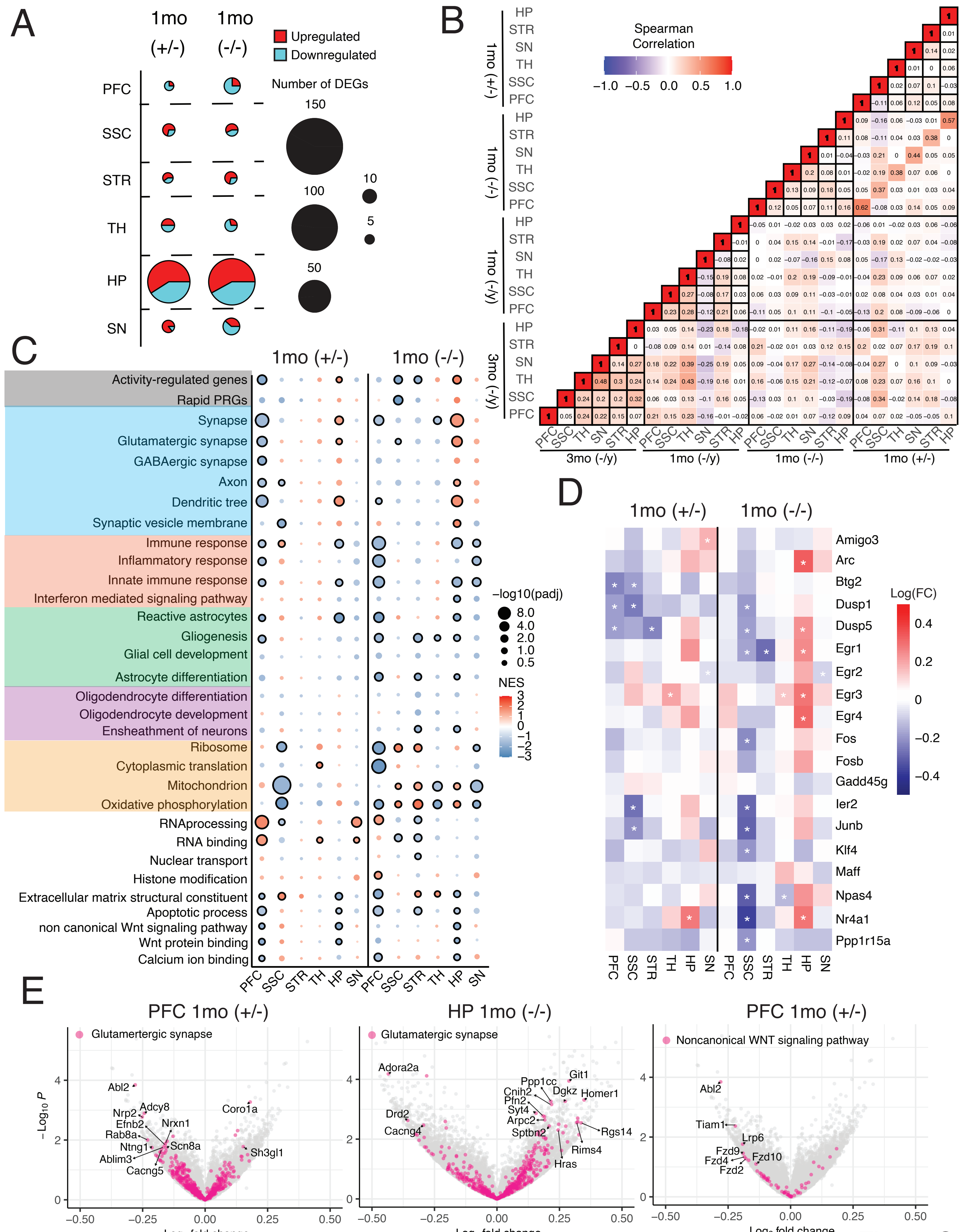

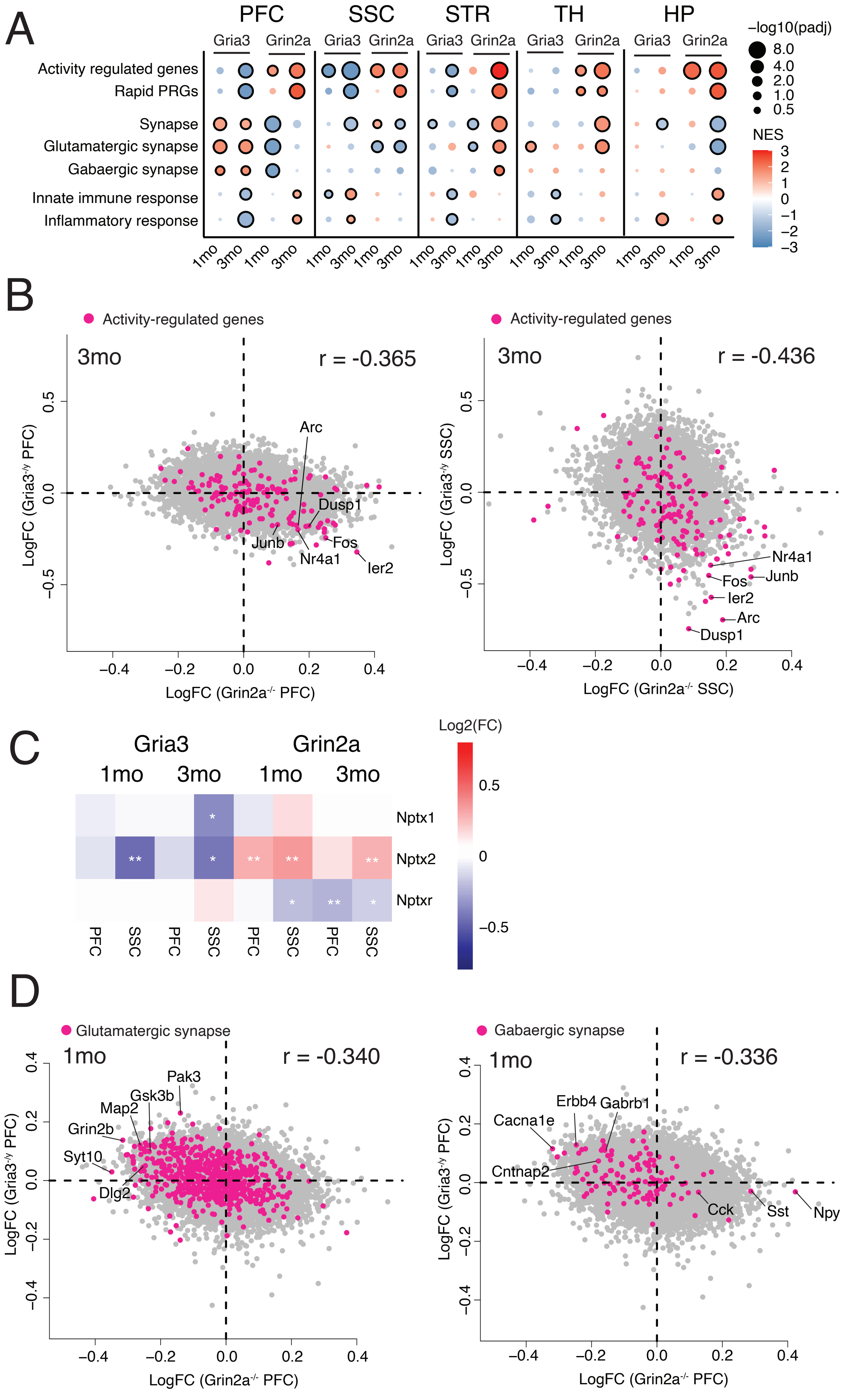
